# Larval development habitats of *Culicoides* midges in the western United States

**DOI:** 10.1101/2025.02.03.636300

**Authors:** Carly Barbera, Christie Mayo, T Alex Perkins, Jason R Rohr

**Affiliations:** Department of Biological Sciences, University of Notre Dame, Notre Dame, IN 46556; Department of Microbiology, Immunology, and Pathology, Colorado State University, Fort Collins, CO 80523

## Abstract

*Culicoides* midges are vectors of bluetongue virus (BTV), an arbovirus affecting wild and domestic ruminants. Bluetongue distribution generally overlaps with vector range, so understanding the vector’s ecology is necessary for predicting BTV risk. *Culicoides* require moist substrate for oviposition and development, and *C. sonorensis*, the primary vector in the western United States, is classically associated with livestock wastewater ponds. However, it is well-known that BTV can be found outside of managed livestock areas, with transmission also occurring in natural settings. To better classify development habitat for *Culicoides*, we conducted a broad biweekly survey of moist and wet habitats from June to September of 2022 in northern Colorado at ten sites, including large livestock operations, non-commercial domestic operations, and natural spaces. Samples were maintained in the insectary for 11 weeks and monitored for emergence of adult midges. Standing water substrates displayed higher presence and abundance of midges than running or transient habitats, though all microhabitats showed emergence. Additionally, livestock sites did not produce more midges than other site types, and in fact more midges were observed in natural spaces per sample and overall. Livestock spaces did, however, show significantly higher proportions of *C. sonorensis* midges, which are thought to be the most competent vectors of BTV in this region. These results suggest that development sites in natural areas may play an important role in maintaining vector populations in the western U.S. outside of previously implicated livestock operations, and that differences in larval habitat lead to differences in vector species composition.

## Background

Midges of the genus *Culicoides* are small, biting insects of veterinary and agricultural importance because of their role as vectors of pathogens affecting ruminants, such as cattle. Among these pathogens are bluetongue virus (BTV) and epizootic hemorrhagic disease virus (EHDV), both of which severely impact wildlife and domesticated populations of ruminants. BTV is present on all continents except for Antarctica, with distributions generally reflecting the range in which competent vectors occur (MacLachlan 2004, Tabachnick 1996). This range has been expanding in recent years, with latitudinal limits now beyond the historical bounds of 40-50 °N and 35-40 °S (Mayo et al. 2020). This range expansion, combined with an increase in outbreak frequency and incursion of exotic serotypes in many regions, has caused severe agroeconomic consequences. Notably, *Culicoides* vectors have also been implicated as competent vectors for Oropouche virus (OROV), which has raised alarm because of an ongoing outbreak in Central and South America (Gallichotte et al. 2024, Mahopatra et al. 2024). Thus, the expansion of these pathogens highlights a need for deeper knowledge of the factors driving transmission of *Culicoides*-borne pathogens.

Immature stages of *Culicoides* are typically found in semiaquatic habitats, such as swamps, pond and stream edges, animal manure, and tree holes. Different species of midges are associated with different larval development habitats, with some displaying more generalist tendencies, and some with a narrower range of preferred habitats (Mullens et al. 2015). In the western US, the primary implicated vector species for BTV and EHDV is *Culicoides sonorensis* (Mayo et al. 2020, McGregor et al. 2022). This species is classically associated with livestock habitats, as wastewater ponds on livestock operations are commonly implicated as breeding grounds (Purse et al. 2015, Erram et al. 2017). However, there are regular cases of BTV and EHDV among wildlife where wastewater ponds do not occur (Roug et al. 2012, Dorak et al. 2022, Fox et al. 2015). There is some evidence that there may be more ephemeral sites for *C. sonorensis* development, such as in urban/industrial ponds and puddles, as well as more natural habitats, such as creeks, waterfowl refuge, and bison wallows (Mullens et al. 2015, Pfannensteil et al. 2015). Additionally, absence of wastewater ponds on livestock operations has not been shown to reduce numbers of adult *Culicoides* (Mayo et al. 2014, Harrup et al. 2014), further indicating that these may not be the sole larval development sites. Without specific knowledge of the range of habitats where *C. sonorensis* can develop, predictions of vector distribution, and in turn transmission, may be unreliable.

It has also been suggested that *C. sonorensis* is not the only vector species responsible for transmission of BTV in the Western United States. While the geographic distribution of BTV correlates with *C. sonorensis* distribution, this does not seem to be the case for EHDV (Tabachnik et al. 1996, Ruder et al 2015), particularly in eastern and central US, where other midge species are suspected, but have not been confirmed, as vectors (Pfannenstiel et al. 2015). In fact, a sole vector species is not typical; many locations have a diversity of species acting as vectors (Purse et al. 2015). Additionally, both Pfannestiel et al. (2015) and Purse et al. (2015) noted that *C. sonorensis* is one of the few species that can be reared in the lab, and thus it is likely to have received disproportionate attention compared to other potential vector species. Indeed, previous molecular work found BTV in other midge species (per. obs.), many of which were collected in non-domestic environments, supporting the idea that alternative environments and alternative vector species can contribute to BTV transmission.

Several studies have noted the need to further describe habitat suitability for *Culicoides* development in areas with ongoing BTV and EHDV transmission (Mullens et al. 2015, Purse et al 2015, Pfannestiel et al 2015). More specifically, there is a lack of work establishing the environments that are conducive to *Culicoides* vector development and persistence, and to what extent these environments contribute to vector populations. To characterize conditions that support *Culicoides* development, we conducted a field survey of potential *Culicoides* immature development habitats across different landscape types, including a range of moist substrates on intensively managed livestock operations, rangeland, mixed-host domestic operations, and natural land. We aimed to test the hypothesis that alternative habitat types, outside of the classically implicated livestock wastewater ponds, are providing significant development habitat for *Culicoides* midges. Additionally, we aimed to establish habitat associations for non-*sonorensis Culicoides* species, which might also have a potential role in BTV and EHDV transmission.

## Materials and Methods

### Sample collection and processing

Ten field sites in northern Colorado were selected representing a range of environments. Different locations will be referred to as “sites,” and “site type” will describe the three land use types of each site: livestock operation describes commercial livestock sites; non-commercial operation describes non-commercial domestic, mixed-host sites; natural space describes open space with non-domestic ruminant hosts. Predictor variables are defined in more detail in Table 1. At each site, possible habitat for larval midges was identified as any visible moist substrate, including naturally occurring ponds and streambeds, wastewater ponds, irrigation canals, hoofprints, and puddles (Figure 1).

**Table 1.** A summary of the predictor variables and their definitions used in all models.

| Predictor Variable | Definition |
| --- | --- |
| *Site Type* |  |
| Livestock Operation | Commercial livestock operations (dairies, feedlots, and range sites containing either cattle or sheep) |
| Non-Commercial Operation | Domestic, mixed-host sites (research facilities, equestrian sites, and small home farm) |
| Natural Space | Open spaces populated primarily by wild host species (state park, wildlife refuge) |
| *Substrate Type* |  |
| Running | Consistently flowing water sources (streams and creeks, drainage ditches) |
| Standing | Non-flowing, consistently present water bodies (agricultural ponds, natural ponds, swampy areas) |
| Transient | Ephemeral water bodies (puddles, temporary overflow from nearby water sources) |

**Figure 1.**
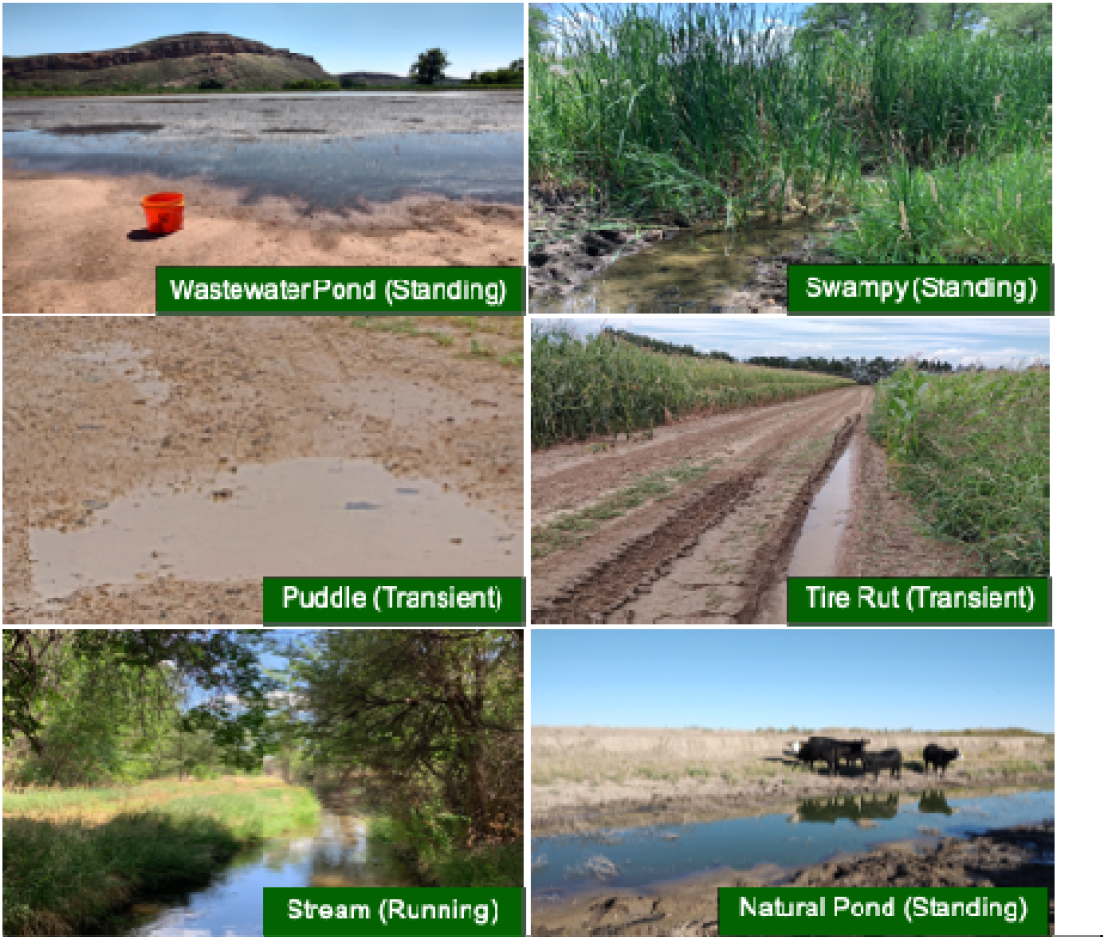
Examples of sampling locations for substrates and their categorizations at different field sites in Colorado, 2022.

“Sample” will refer to a unit of substrate taken from a given sampling location from a given site on a given sampling day. Each sample is classified as one of three ‘‘substrate types:” running, standing, or transient (Table 1). Most sampling locations were visited more than once throughout the season, but some were not, depending on whether the substrate at the location was still available at different sampling dates. All available identified substrate types were sampled at each site visit.

Substrate collection occurred between late June and early September, 2022. Samples were taken using a trowel to skim the edge of the sampling location, focusing on the edges of the substrate body, where the mud or ground met the water body. Samples were placed in sealed plastic containers and transported to the insectary that day, where 150 g of substrate was transferred from each sample to a sealed container in the lab with organza covering to allow oxygen transport but prevent escape of emerging midges. The insectary was kept on a 12 hr light cycle, at ∼30 °C and 60-80% relative humidity. Moisture level of each sample was maintained by adding water as needed to the original mass of 150 g. Each sample was monitored for emergence of adults over the course of 11 weeks, after which we disposed of substrate samples. Emerging adults were captured with a small portable insect aspirator. Adults were then counted and visually identified, when possible, under the microscope as *Culicoides sonorensis* or other *Culicoides* species.

### Statistical analysis

All statistical analyses were performed in ‘Rstudio’ software version 2024.4.2 (R Core Team 2024). We used generalized linear mixed effects models in package ‘glmmTMB’ (Brooks et al. 2017) to determine effects of site type and substrate type on midge emergence. Models with an interaction between site type and substrate were tested and rejected due to higher AIC values than models excluding the interaction. We ran separate models for each of four response variables: ‘Abundance (zeros included),’ which refers to the total number of midges emerged from each sample, including those without any midges (n = 196); ‘Presence,’ which refers to the proportion of overall samples that had any adult midges emerge over the course of 11 weeks in the insectary (n = 196); ‘Abundance (zeros excluded),’ which refers to the total number of midges emerged from only those samples containing midges (n = 57); and ‘Proportion *C. sonorensis*,’ which refers to the proportion of emerged midges in each sample that were visually identified as *Culicoides sonorensis* (n = 54, some samples unidentifiable). Non-commercial operations were excluded from the proportion *C. sonorensis* model because there was only one sample containing midges within this category, and thus the category had no detectable variance. We used a binomial distribution for both presence and proportion *C. sonorensis*, and a negative binomial distribution for abundance models. All models included site type and substrate as categorical fixed effects. Each model also included a random effect of sampling week, to account for changes in larval abundance over the course of the season. The abundance (zeros included) and presence models included a random effect of sampling location nested within site, to account for the non-independence of repeated sampling at each location.

For models with significant effects of fixed predictors, we used the ‘emmeans’ package version 1.10.1 (Lenth 2024) on each of the models to conduct post-hoc analysis, testing for pairwise differences between treatments within a factor.

## Results

### Data

A total of 196 samples were collected, 57 of which had adult midges emerge, for a total of 688 emerged midges. Midges emerged from samples as early as one day after collection to up to 61 days after collection.

### Effect of Site Type

Site type was not a significant predictor of total abundance of midges across all samples, including those with no midges (p = 0.145, R^2^ = 0.641; Figure 2A, Table 2A) or the proportion of samples with midges present (p = 0.167, R^2^ = 0.465; Figure 2B, Table 2B). Though differences between groups were not significant, natural spaces did display higher abundance (mean predicted value = 2.61, CI = 0.41 - 16.52) than both livestock operations (mean predicted value = 0.56, CI = 0.18 - 1.74) and non-commercial operations (mean predicted value = 0.27, CI = 0.03 - 2.60) (Figure 2A, Table 2A). In the model excluding samples with no midges present, site type also was not a significant predictor of midge abundance (p = 0.082, R^2^ = 0.550; Figure 2C, Table 2C).

**Figure 2.**
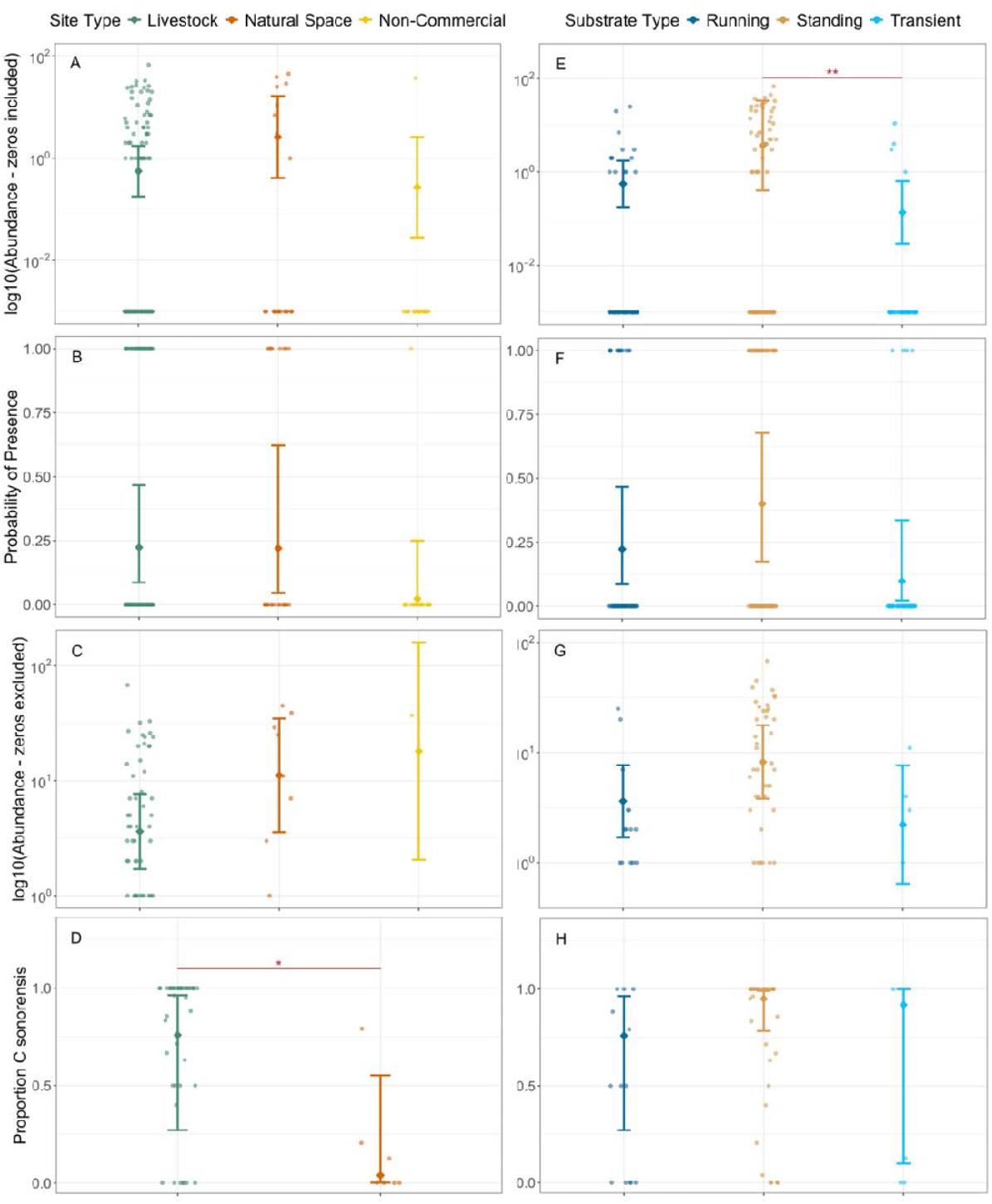
Predicted effects of site type (panels A - D) and substrate type (panels E - H) on the abundance of midges emerging from samples, zeros included (panels A and E), the proportion of samples with midges present (panels B and F), the abundance of midges emerging from samples, zeros excluded (panels C and G), and the proportion of midges per sample that are identified as *C. sonorensis* (panels D and H). Dots show mean predictions per category, and bars around them show the 95% confidence intervals.

**Table 2.**
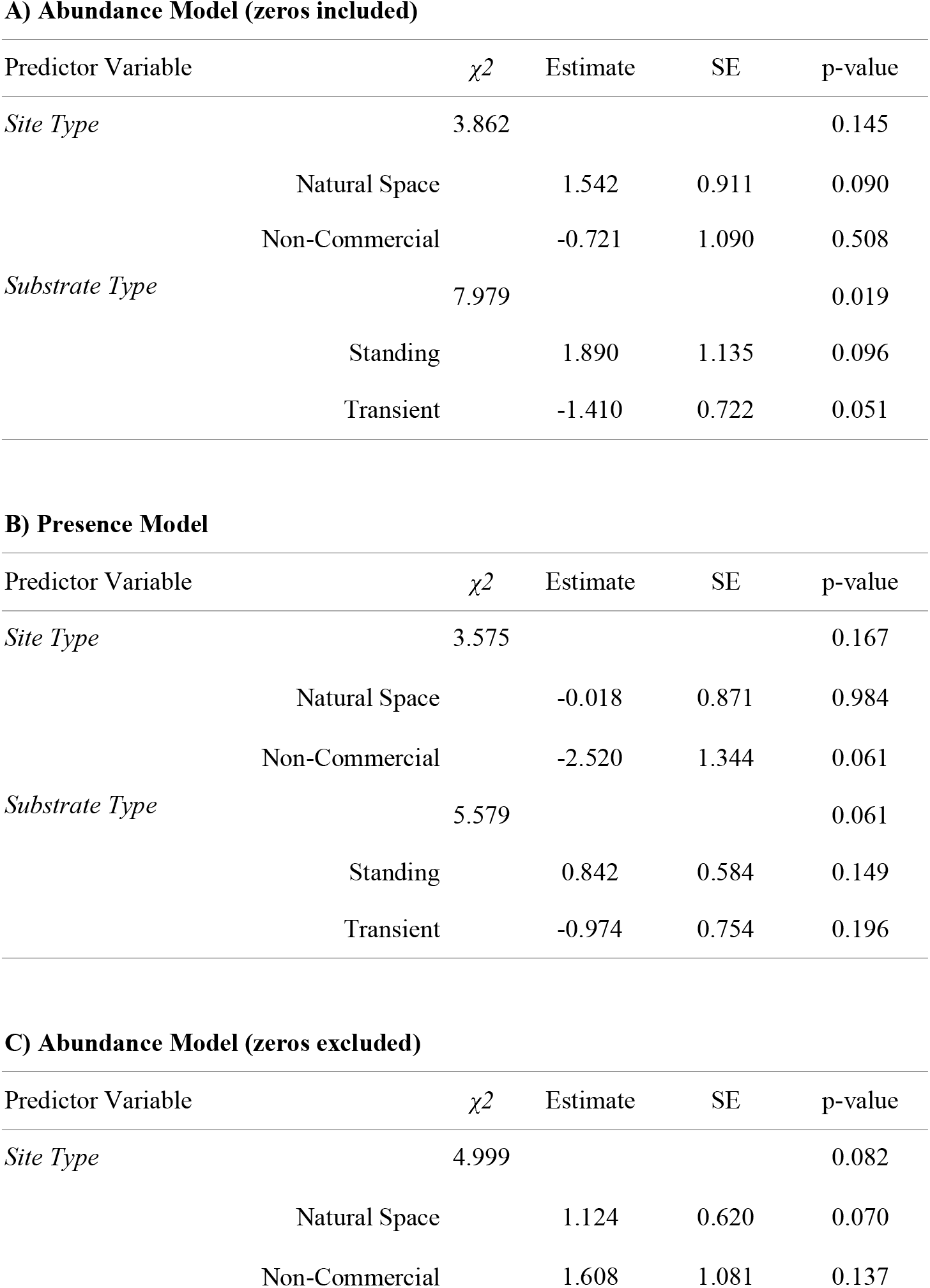
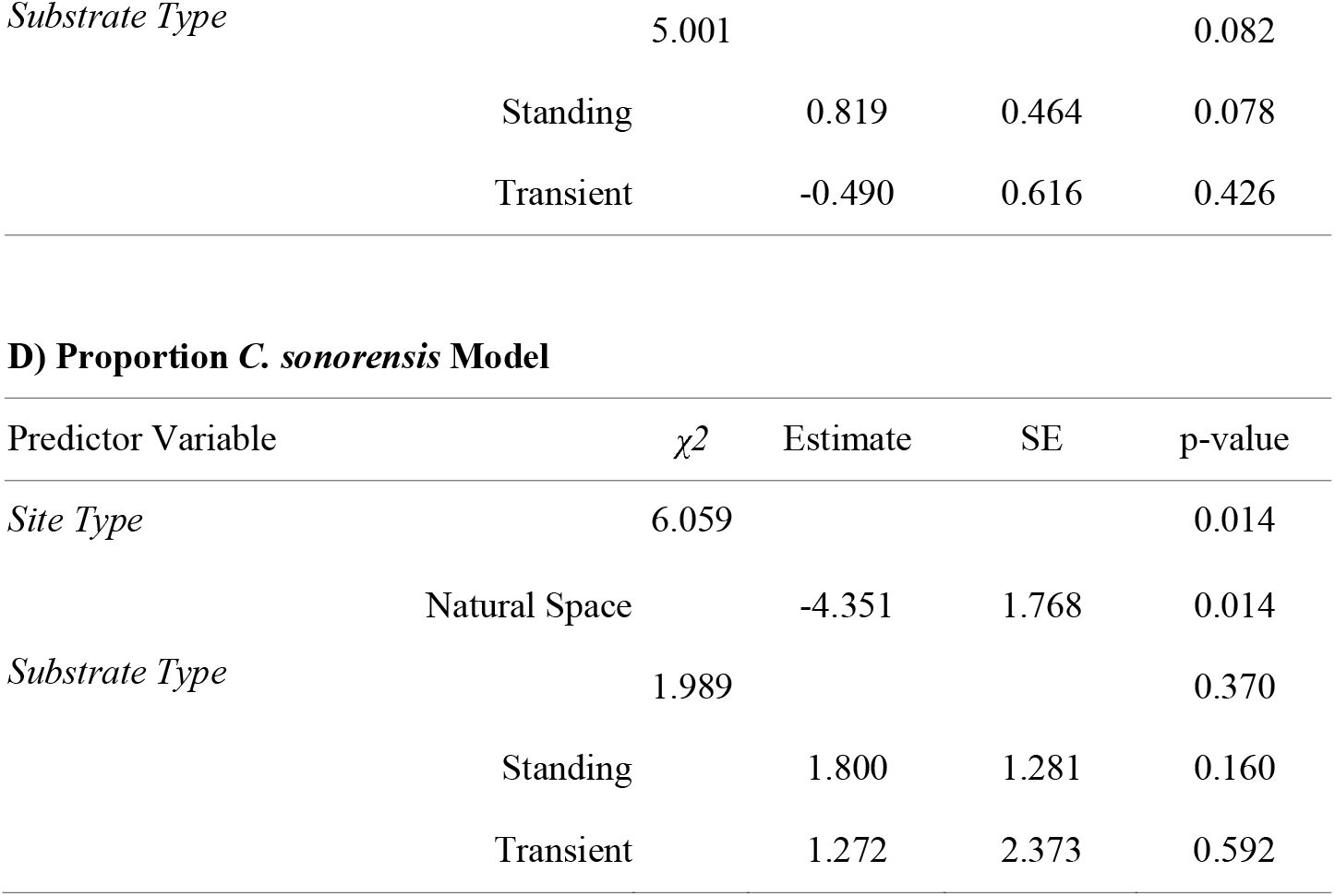
Summary statistics for the three models in the analysis. A) GLMM with the total number of midges emerged from samples, even those without midges, over the course of 11 weeks in the insectary as the response variable (Abundance, zeros included), using a negative binomial distribution (n = 196). B) GLMM with proportion of samples containing any midges as the response variable (Presence), using a binomial distribution (n=196). C) GLMM with the total number of midges emerged from samples containing midges as the response variable (Abundance, zeros excluded), using a negative binomial distribution (n = 57). D) GLMM with proportion of midges in each sample that were visually identified as *Culicoides sonorensis* as the response variable (Proportion *C. sonorensis*), using a binomial distribution and weighted by the number of midges in that sample that had been id’ed, i.e., the sample size for that particular sample (n = 55).

Site type was a significant predictor of the proportion of *C. sonorensis* (p = 0.014, R^2^ = 0.963), with livestock operations (mean predicted value = 0.76, CI = 0.27 - 0.96) showing a significantly higher proportion of *C. sonorensis* than natural spaces (mean predicted value = 0.04, CI = 0.00 - 0.55) (Figure 2D, Table 2D).

### Effect of Substrate

Substrate type was a significant predictor of abundance of midges emerging from samples, including those without midges (p = 0.019). In particular, a pairwise comparison showed significantly more emergence from standing substrate (mean predicted value = 3.69, CI = 0.40 - 33.70) than from transient substrate (mean predicted value = 0.14, CI = 0.03 - 0.63), though there was no significant difference between running water (mean predicted value = 0.56, CI = 0.18 - 1.74) and other groups (Figure 2E, Table 2A). Presence of midges in samples was marginally not affected by substrate type (p = 0.061, Figure 2F, Table 2B). When samples without midges were excluded, substrate type was not a significant predictor of midge abundance (p = 0.082; Table 2G, Figure 2C). Substrate type was also not a significant predictor of proportion of *C. sonorensis* (p = 0.370; Figure 2H, Table 2D).

## Discussion

The results from this study demonstrate that *Culicoides* midges are able to emerge from variable landscapes and microhabitat types, and are not limited to classically implicated livestock environments for development. Perhaps most striking is that natural spaces may be contributing to *Culicoides* midge emergence as much as or more than livestock environments. Though the proportion of samples with midges present appeared to be similar between livestock areas and natural spaces, the number of midges emerging from natural spaces overall was higher (Figure 2 A,B, Table 2 A,B). Interestingly, though only one sample from a non-commercial site had any midges at all, that sample had a high abundance of midges, meaning that the predicted abundance of midges on non-commercial sites from the zeros-excluded model was high (Figure 2C) compared to the zeros-included model (Figure 2A). This suggests a stochasticity in midge abundance through time, with high-density pockets sometimes occurring even in areas that do not seem to have consistent midge populations, although we did have a relatively low number of samples from each location to demonstrate this. Perhaps intuitively, standing water produced the highest proportion of samples with midges, and also the most midges in those samples (Figure 2E-G). However, running water and transient water also produced midges, and thus cannot be ruled out when considering sources of midge populations. Further, the summer that these data were collected had low overall precipitation in the region compared to other years (Scott 2023). Hence, it is likely that transient substrate was less available as a breeding ground during this study, and could play a larger role in other years.

According to our data, *C. sonorensis* is associated with livestock habitats (Fig 2D, Table 2D). *C. sonorensis* is the primary recognized vector for BTV and EHDV in this region, but the presence of other *Culicoides* midges emerging in places where these pathogens are known to circulate among non-livestock species highlights the need to explore the role the other species play in transmission. Earlier collections of adult midges on these sites showed that non-*sonorensis* midges were primarily *C. crepuscularis*, and qRT-PCR analysis detected BTV in these samples. These findings are supported by other studies that found pools of field-collected *C. crepuscularis* to be positive for BTV (Becker et al 2010, Becker et al 2020, White et al. 2005), despite being associated with avian feeding (Sloyer et al. 2019). Given the lack of data on *C. crepuscularis* and its competence as a vector, it is difficult to know how large of a role the species plays in maintaining transmission. As the dominant species in non-commercial landscapes, the level of competence of *C. crepuscularis* has implications for how effectively transmission of BTV can be maintained among wildlife populations, and in turn, how much of a threat *C. crepuscularis* poses when overlapping with livestock populations.

In addition to giving insight into BTV and EHDV dynamics, this work is important in the context of the current Oropouche virus epidemic in the Americas. The recent surge in cases and expansion into non-endemic areas has highlighted the need for a greater understanding of the ecology of the virus. While the virus is not currently circulating in the United States, *Culicoides* midges, in particular *Culicoides paraensis*, have been identified as instrumental vectors for transmission to humans (Mohapatra et al. 2024). This species is present in North and South America. Additionally, *C. sonorensis* has been shown to be an efficient vector in the lab (Guagliardo et al. 2024, Gallichotte et al. 2024), though it is unclear how well this species can transmit under field conditions (McGregor et al. 2021). Given that multiple species of *Culicoides* are competent vectors of OROV, characterizing the larval ecology of different *Culicoides* species is important to prepare for the possibility that the virus expands to the US.

Our findings may also have implications for vector control. In particular, strategies targeting immature populations are complicated by the presence of many possible microhabitats across the landscape. If large, accessible wastewater ponds were the dominant breeding grounds, then it would be more straightforward to target these areas for larvicide application. However, alternate breeding grounds could maintain adult populations even after wastewater sources are targeted, as well as being more difficult and unpredictable to find and treat. The idea that solely targeting wastewater ponds may not be effective is supported by previous work that found that experimentally removing wastewater ponds on farms did not significantly reduce *Culicoides* abundance when compared to unmodified farms (Mayo et al. 2014, Harrup et al. 2014).

While this study has given insight into vector dynamics in this system, there were some limitations to our approach, and future work is needed to expand upon these results. Although our data suggest that a variety of habitats can support midge populations, larger sample sizes and wider sampling efforts will assist in elucidating the relative contribution of each habitat type to midge population densities. Additionally, surveying across years, to capture changes in habitat availability in years where there is more moisture present, will give more information about the potential for different locations to support larvae. More rigorous sampling effort will also help to capture natural temporal and spatial stochasticity in midge emergence to better quantify how this stochasticity affects population estimates. Furthermore, given our findings of non-*sonorensis* (likely *C. crepuscularis)* midges, especially as they are associated with different habitats from *C. sonorensis*, more work is needed to establish the role that alternative *Culicoides* species play in the transmission of BTV and EHDV. In particular, more insight into the extent that adult populations are associated with different landscapes, whether different vector species display different host feeding preferences, and how efficient these vectors are at transmitting pathogens are needed to understand the role of various vector species in maintaining transmission.

Here, we confirmed the importance of non-domestic landscapes in supporting *Culicoides* populations, and highlighted the need for more work to understand how different species and habitats contribute to disease dynamics. Our findings broaden understanding of *Culicoides* ecology and thus contribute to a greater understanding of BTV and EHDV transmission and potential interventions. Additionally, our work helps to broaden our knowledge of *Culicoides* ecology generally, which is particularly important in light of the recent Oropouche epidemic.

## Acknowledgements

This work was funded by USDA-NIFA AFRI grant #2019-67015-28982 as part of the joint USDA-NSF-NIH-BBSRC-BSF Ecology and Evolution of Infectious Diseases program, and USDA-NIFA grant #2021-38420-34065, NSF grant #DEB-2109293, ITE-2333795, BCS-2307944, the Frontier Research Foundation, and the Alliance Bioversity-CIAT. We gratefully acknowledge producers, staff, and herd managers of the farms from which the samples were obtained. We also thank the research personnel who made this work possible, including Katie James, Colin Korte, Aubrie McGinty, Leah Ortowski, and Tim Wolbers.

## Author’s contributions

CB, CM, TAP, and JRR conceived the study, and CB designed the study. CB collected and processed the data, and CM provided access to insectary facilities and field vehicles. CB analyzed the data and JRR provided guidance on analyses. CB drafted the manuscript, and all authors contributed feedback and approved the final version.

## Notes

### Competing Interest Statement

The authors have declared no competing interest.

## References

1. Becker M, Reeves W, Dejean S, Emery M, Ostlund E, Foil L. Detection of bluetongue virus RNA in field-collected Culicoides spp.(Diptera: Ceratopogonidae) following the discovery of bluetongue virus serotype 1 in white-tailed deer and cattle in Louisiana. Journal of medical entomology. 2014;47(2):269–73.

2. Becker ME, Roberts J, Schroeder ME, Gentry G, Foil LD. Prospective study of epizootic hemorrhagic disease virus and bluetongue virus transmission in captive ruminants. Journal of medical entomology. 2020;57(4):1277–85.

3. Brooks ME, Kristensen K, Van Benthem KJ, Magnusson A, Berg CW, Nielsen A, et al. glmmTMB balances speed and flexibility among packages for zero-inflated generalized linear mixed modeling. The R journal. 2017;9(2):378–400.

4. Dorak SJ, Varga C, Ruder MG, Gronemeyer P, Rivera NA, Dufford DR, et al. Spatial epidemiology of hemorrhagic disease in Illinois wild white-tailed deer. Scientific reports. 2022;12(1):6888.

5. Erram D, Zurek L. Larval Development of Culicoides sonorensis (Diptera: Ceratopogonidae) in Mud Supplemented With Manure of Various Farm Animals. Journal of Medical Entomology. 2018;55(1):43–50.

6. Fox KA, Diamond B, Sun F, Clavijo A, Sneed L, Kitchen DN, et al. Testicular lesions and antler abnormalities in Colorado, USA mule deer (Odocoileus hemionus): a possible role for epizootic hemorrhagic disease virus. Journal of Wildlife Diseases. 2015;51(1):166–76.

7. Gallichotte EN, Ebel G, Carlson CJ. Vector competence for Oropouche virus: a systematic review of pre-2024 experiments. medRxiv. 2024:2024.10. 17.24315699.

8. Guagliardo SAJ, Connelly CR, Lyons S, Martin SW, Sutter R, Hughes HR, et al. Reemergence of Oropouche Virus in the Americas and Risk for Spread in the United States and Its Territories, 2024. Emerging Infectious Diseases. 2024;30(11):2241.

9. Harrup L, Gubbins S, Barber J, Denison E, Mellor P, Purse B, et al. Does covering of farm-associated Culicoides larval habitat reduce adult populations in the United Kingdom? Veterinary parasitology. 2014;201(1-2):137–45.

10. Lenth R. Estimated Marginal Means, Aka Least-Squares Means [R Package Emmeans Version 1.10. 1]. R-project org. 2024.

11. Maclachlan NJ. Bluetongue: history, global epidemiology, and pathogenesis. Preventive veterinary medicine. 2011;102(2):107–11.

12. Mayo C, McDermott E, Kopanke J, Stenglein M, Lee J, Mathiason C, et al. Ecological dynamics impacting bluetongue virus transmission in North America. Frontiers in Veterinary Science. 2020;7:186.

13. Mayo CE, Osborne CJ, Mullens BA, Gerry AC, Gardner IA, Reisen WK, et al. Seasonal variation and impact of waste-water lagoons as larval habitat on the population dynamics of Culicoides sonorensis (Diptera: Ceratpogonidae) at two dairy farms in northern California. PLoS One. 2014;9(2):e89633.

14. McGregor BL, Connelly CR, Kenney JL. Infection, dissemination, and transmission potential of North American Culex quinquefasciatus, Culex tarsalis, and Culicoides sonorensis for Oropouche virus. Viruses. 2021;13(2):226.

15. McGregor BL, Shults PT, McDermott EG. A review of the vector status of North American Culicoides (Diptera: Ceratopogonidae) for bluetongue virus, epizootic hemorrhagic disease virus, and other arboviruses of concern. Current Tropical Medicine Reports. 2022;9(4):130–9.

16. Mohapatra RK, Mishra S, Satapathy P, Kandi V, Tuglo LS. Surging Oropouche virus (OROV) cases in the Americas: A public health challenge. New Microbes and New Infections. 2024;59.

17. Mullens BA, McDermott EG, Gerry AC. Progress and knowledge gaps in Culicoides ecology and control. 2015.

18. Pfannenstiel RS, Mullens BA, Ruder MG, Zurek L, Cohnstaedt LW, Nayduch D. Management of North American Culicoides biting midges: current knowledge and research needs. Vector-borne and zoonotic diseases. 2015;15(6):374–84.

19. Pfannenstiel RS, Ruder MG. Colonization of bison (Bison bison) wallows in a tallgrass prairie by Culicoides spp (Diptera: Ceratopogonidae). Journal of Vector Ecology. 2015;40(1):187–90.

20. Purse B, Carpenter S, Venter G, Bellis G, Mullens B. Bionomics of temperate and tropical Culicoides midges: knowledge gaps and consequences for transmission of Culicoides-borne viruses. Annual Review of Entomology. 2015;60(1):373–92.

21. Roug A, Swift P, Torres S, Jones K, Johnson CK. Serosurveillance for livestock pathogens in free-ranging mule deer (Odocoileus hemionus). PLoS One. 2012;7(11):e50600.

22. Ruder MG, Lysyk TJ, Stallknecht DE, Foil LD, Johnson DJ, Chase CC, et al. Transmission and epidemiology of bluetongue and epizootic hemorrhagic disease in North America: current perspectives, research gaps, and future directions. Vector-Borne and Zoonotic Diseases. 2015;15(6):348–63.

23. Scott W, Jr. Colorado Climate Center 2023 [Available from: https://climate.colostate.edu/co_cag/cag_time.html.]

24. Sloyer KE, Acevedo C, Runkel AE, Burkett-Cadena ND. Host associations of biting midges (Diptera: Ceratopogonidae: Culicoides) near sentinel chicken surveillance locations in Florida, USA. Journal of the American Mosquito Control Association. 2019;35(3):200–6.

25. Tabachnick WJ. Culicoides variipennis and bluetongue-virus epidemiology in the United States. Annual review of entomology. 1996;41(1):23–43.

26. Team RC. R: A language and environment for statistical computing. Foundation for Statistical Computing, Vienna, Austria. 2024.

27. White DM, Wilson WC, Blair CD, Beaty BJ. Studies on overwintering of bluetongue viruses in insects. Journal of General Virology. 2005;86(2):453–62.

